# Investigating Deep Feedforward Neural Networks for Classification of Transposon-Derived piRNAs

**DOI:** 10.1101/2020.04.08.032755

**Authors:** Alisson Hayasi da Costa, Renato Augusto C. dos Santos, Ricardo Cerri

## Abstract

PIWI-Interacting RNAs (piRNAs) form an important class of non-coding RNAs that play a key role in the genome integrity through the silencing of transposable elements. However, despite their importance and the large application of deep learning in computational biology for classification tasks, there are few studies of deep learning and neural networks for piRNAs prediction. Therefore, this paper presents an investigation on deep feedforward networks models for classification of transposon-derived piRNAs. We analyze and compare the results of the neural networks in different hyperparameters choices, such as number of layers, activation functions and optimizers, clarifying the advantages and disadvantages of each configuration. From this analysis, we propose a model for human piRNAs classification and compare our method with the state-of-the-art deep neural network for piRNA prediction in the literature and also traditional machine learning algorithms, such as Support Vector Machines and Random Forests, showing that our model has achieved a great performance with an F-measure value of 0.872, outperforming the state-of-the-art method in the literature.

## 1. Introduction

PIWI-interacting RNAs (piRNAs) comprise a class of small non-coding RNAs (ncRNAs) of approximately 24–31 nucleotides (although this range may change across different species) [1, 2] that are present in animals ranging from sponges to humans [1, 3], and are mainly expressed in gonads [1, 4, 5, 6].

A well-known role of piRNAs is silencing of transposable elements (TEs) in the germline cells similar to other RNA-based mechanisms such as microRNAs (miRNAs) and small interfering RNAs (siRNAs) [1, 3, 4, 5, 6, 7]. In brief, after maturation, piRNAs bound with PIWI proteins – a germline-specific sub-clade of the Argonaute family [1] – to form piRNA-induced silencing complexes (piRISC) that can recognize and silence complementary RNA targets at both the transcriptional and post-transcriptional levels [3, 1, 4, 5].

Although TEs have a significant role in evolution, their mobility in the genome can generate deleterious mutations leading to biological problems, such as infertility [4, 7]. Therefore, silencing of TEs by piRNAs is indispensable to protect the integrity of genomes in germline cells against harmful transposons [4, 5, 8], especially, in animals that undergo obligate sexual reproduction, making this class of small ncRNAs guardians of the genome [4].

The importance of piRNAs brings out the need for efficient identification methods, capable of distinguishing the piRNA sequences from other ncRNAs. However, the development of computational tools for this task is complex [9]. Despite the genomic locations of piRNA clusters being often conserved between related species (such as mouse and humans), piRNA sequences are immensely diverse and rarely conserved [10, 2, 11]. For example, as presented by Weick and Miska [12], in both *Drosophila melanogaster* and vertebrates, mature piRNAs are slightly longer than miRNAs and siRNAs, with sequences between 24 and 31 nucleotides in length, have a preference for a 5’ uracil, and possess a 3’-most sugar that is 2’-O-methylated. On the other hand, *Caenorhabditis elegans* piRNAs are 21 nt long but share the 5’ and 3’ features of piRNAs in other organisms. Therefore, due to the large diversity of piRNA sequences, developing computational methods based on common structural-sequence features among species is challenging [13], making the use of advanced machine learning (ML) approaches, as deep learning (DL), very attractive [14].

Deep learning is now one of the most active fields in machine learning and has been successfully performed many complex tasks such as image and speech recognition, natural language understanding, and most recently, tasks related to computational biology [15]. In the latter, deep learning models are attractive to computational biologists, mainly due to the ability to learn a robust representation directly from raw input data, including bases of DNA sequences or pixel intensities of a microscopy image. On the other hand, traditional machine learning methods require extensive laboratory work for feature extraction in order to develop a reliable model [16].

Since 2016, many studies and methods based on deep learning for solving computational biology problems have been published [16], such as DeeperBind [17], DeepSEA [18], DanQ [19] and many others [16, 15]. However, there are still few studies about the application of deep learning approaches in piRNA prediction [16, 14]. The predominant majority is based on traditional ML models, like Support Vector Machines (SVMs). Some examples are Piano [20], Pibomd [21], piRPred [22], IpiRId [23] and piRNAPredictor of Luo et. al. [13]. But with the exception of piRNAPredictor, all other methods have limitations of use, which include: need for genomic and epigenomic information, can only be applied to specific organisms or have performance problems in different datasets. [14].

The first and only deep learning predictor for piRNAs is called piRNN [14], which is a simple Convolutional Neural Network (CNN). CNN has become very popular in Computer Vision (CV), reaching the state-of-the-art in tasks such as classification and image recognition [24, 25]. However, it still have problems, in particular, high computational costs and slow learning, hard hyperparameter tuning, and the need of large amounts of data [26, 27]. In addition, other models like Deep Belief Networks, or even simple Feedforward Networks (also called Multilayer Perceptrons) [28], can achieve performances comparable to CNNs in other domains than Computer Vision with less computational cost [26, 27].

Considering the lack in the literature regarding applications of deep neural network models in piRNAs prediction, this paper presents an investigation on deep feedforward networks models for classification of transposon-derived piRNAs. We analyze and compare the results of the deep neural networks in different hyperparameters choices, such as number of layers, activation functions and optimizers, clarifying the advantages and disadvantages of each configuration. From this analysis, we propose a model to predict if a sequence is a human transposon-derived piRNA or not. Furthermore, we compare our method with the state-of-the-art deep neural network for piRNA prediction in the literature and also traditional machine learning algorithms, such as Support Vector Machines and Random Forests. Additionally, it is expected that the analyzes and comments encourage the application of DFNs in other classes of small ncRNAs.

The remainder of this article is organized as follows. Section 2 explains the methodology used for the dataset acquisition, process of extracting features from sequences, preprocessing algorithms applied to data, network architectures, hyperparameters chosen, and the process of training and testing for hyperparameter optimization and comparison with other methods. Section 3 presents the results obtained by the DFNs, analyzing the impact of each hyperparameter choice in the final result. Moreover, we compare our proposal with other traditional machine learning algorithms and piRNN. Finally, Section 4 presents the conclusions obtained in this work, and future research directions.

## 2. Methodology

### 2.1. Datasets and Sequence Feature Extraction

The dataset used for the experiments has a total of 14,810 samples, where 7,405 are transposon-derived piRNA sequences (positive samples) and 7,405 are pseudo-piRNA sequences (negative samples). All samples are *Human* ncRNAs and were obtained from the supplementary material provided in the work of Luo et. al. [13], where it is possible to find details about its construction. Figure 1 shows the sequence lengths of positive and negative samples.

**Figure 1:**
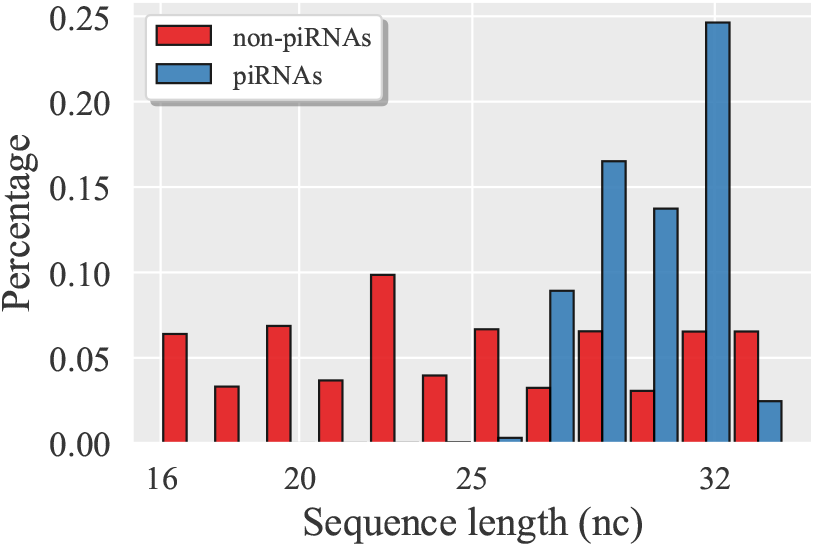
Length distribution of piRNAs and non-piRNAs sequences.

From the data collected, three different feature sets were extracted using the Pse-in-One-2.0 tool (local version) [29]. Spectrum profile, also named *k*-mer feature, counts the occurrences of *k*-mer motif frequencies (*k*-length contiguous strings) in sequences. Mismatch profile also counts the occurrences of *k*-mers, but allows max *m* (*m ≤ k*) inexact matching, which is the penalization of spectrum profile. Subsequence profile considers not only the contiguous *k*-mers but also the non-contiguous *k*-mers, and the penalty factor *w* (0 *≤ w ≤* 1) is used to penalize the gap of non-contiguous *k*-mers [13]. Furthermore, since features have parameters that can be adjusted (*k*, *m* and *w*, where *k* is present in all three features), for *k*-mers we adopted *k* = 1, 2, 3, 4. For mismatch profile we used (*k*, *m*) = (1,0), (2,1), (3,1), (4,1) and for subsequence profile (*k*,*w*) = (1,1), (2,1), (3,1), (4,1). Thus, a total of 340 attributes per sequence (sum of 4^1^, 4^2^, 4^3^, 4^4^, where *k* is the exponent) was obtained in each of the feature sets. After the feature extraction process, two feature scaling algorithms were applied on each of the obtained feature sets.

The normalization (also known as Min-Max Normalization), which scales and translates each feature individually such that it is in a range of 0 to 1; and the standardization (also known as Z-Score Normalization), which transforms the original data distribution into a normal distribution with zero mean and unit variance. Therefore, a total of nine datasets with 340 attributes each was obtained. Three corresponding to raw *k*-mers, mismatch and subsequence (i.e, before feature scaling), three corresponding to these same features, but after Min-Max Normalization, and three others after Z-Score Normalization.

### 2.2. Architecture and Hyperparameters

We implemented a Deep Feedforward Network with eight different hyperparameter configurations, where each one is characterized by the number of hidden layers, activation function and optimizer used. As for the number of hidden layers, variations of three and five hidden layers were implemented with 340 units per layer (number equivalent to the input array dimension), and a dropout layer with 0.5 dropout ratio between all layers [30]. These numbers were chosen to verify how increasing depth can improve or impair the generalization capacity of the model, together with the activation function and the optimizer used [31, 32, 33]. In the output layer, a single neuron with sigmoid was used to predict if a sequence is a transposon-derived piRNA or not [32].

The activation functions selected were logistic function (sigmoid) and Rectified Linear Unit (ReLU). Although the sigmoid is very efficient to deal with sparse data, when used in a neural net with a large number of layers, the gradients may become vanishingly small, preventing learning from occurring [32, 31]. On the other hand, ReLU is the activation state-of-the-art and has important qualities, such as sparse activation and better gradient propagation, allowing neural networks with a large number of hidden layers with less vanishing gradient. However, the sparse activation along with the large natural sparsity of data can cause a accumulation of large error gradient values, resulting in large updates to the network weights and, consequently, an very unstable model [34, 35]. Thus, given the pros and cons of each activation function, we analyzed the efficiency of both in the classification of piRNAs.

Moreover, we used the Glorot weight initialization [31] in layers whose activation was sigmoid to minimize the occurrence of vanishing gradient, and He weight initialization [34] in the layers with ReLU to promote a faster and efficient convergence, in addition to dealing with vanishing and exploding gradient problems.

The adopted optimizers were the Stochastic Gradient Descent (SGD) [28], with a learning rate of 0.01 and Nesterov Momentum of 0.9, and the Adaptive Moment Estimation (Adam), with default parameters provided in its original paper [36]. Both were chosen in order to verify their ability to produce a good generalization and dealing with possible difficulties of poor hyperparameters choice. Finally, the cost function used was the log loss.

All implementations and experiments were performed using Python 3.6.5, Theano 1.0.3 [37], Keras 2.2.0 [38], and scikit-learn 0.20.1 [39]. Table 1 presents the eight hyperparameter configurations implemented, where the X in DFNX stands for the number (identification) of the experiment performed.

**Table 1:** The eight DFN configurations implemented

| **Model** | **Layers** | **Activation** | **Optimizer** |
| --- | --- | --- | --- |
| DFN1 | 3 | ReLU | Adam |
| DFN2 | 3 | ReLU | SGD |
| DFN3 | 3 | sigmoid | Adam |
| DFN4 | 3 | sigmoid | SGD |
| DFN5 | 5 | ReLU | Adam |
| DFN6 | 5 | ReLU | SGD |
| DFN7 | 5 | sigmoid | Adam |
| DFN8 | 5 | sigmoid | SGD |

### 2.3. Evaluation Metrics

We used three evaluation metrics to assess the performance of the models: Recall (REC) (Equation 2), Precision (PRE) (Equation 1) and F1-Measure (F) (Equation 3). In these equations, *tp*, *fp* and *fn* stands, respectively, for the number true positive samples, the number of false positive samples, and the number of false negative samples.

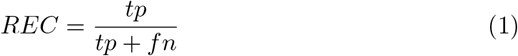

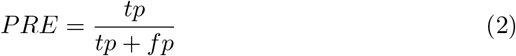

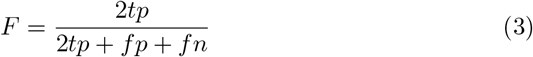

In order to better analyze the results of each hyperparameter configuration, and then perform a comparison with the state-of-the-art piRNN literature method, each one of the nine datasets was splitted into two disjoint subsets, G1 and G2. The G1 group was used for hyperparameter tuning, and the G2 group was used in the comparison with piRNN and other traditional machine learning algorithms, allowing a comparison with the least possible bias. Thus, the subsets of group G1 have a total of 7,406 samples divided into 3,703 positive samples and 3,703 negative samples, and the subsets of group G2 have 3,702 positive samples and 3,702 negative samples.

In the hyperparameter optimization step, all models were trained and evaluated using the 10-Fold Cross-Validation strategy, with 256 epochs and a batch size of 32. The results obtained by the models in the test fold were used to analyze, evaluate and compare the impact of different hyperparameter configurations. Then, the best performing model in the hyperparameter selection step was trained and tested also using 10-Fold Cross-Validation, 256 epochs and a batch size of 32. However, we now used the samples of group G2.

## 3. Results and Discussion

To well understand the results, we must first understand the data used since raw data and preprocessed data (i.e, after applying feature scaling) affect the results in different ways. For example, *k*-mers count the occurrences of *k*-mer motif frequencies in sequences, so the values are limited to a range of 0 to 1, making the application of Min-Max Scaling not as efficient as it could be. However, the same is not true for mismatch profile and subsequence profile.

Both features (mismatch and subsequence) in their raw state are composed of positive integers, with mismatch ranging from 0 to 30 and subsequence ranging from 0 to 14950. These very sparse interval not only express the diversity of input samples but also the presence of several outliers. In addition, since the piRNA sequences are small in length, the higher the value of *k*, the lower the frequency. Consequently, the first 84 attributes (sum of 4^1^, 4^2^, 4^3^) have a slightly better behavior than the other 256 attributes (equivalent to 4^4^), as shown in Figures 2 and 3. Therefore, the application of Min-Max Normalization and Z-Score Normalization (although sensitive to outliers) is extremely useful and necessary for these features by reducing the interval to which they belong and facilitating the convergence of the neural network.

**Figure 2:**
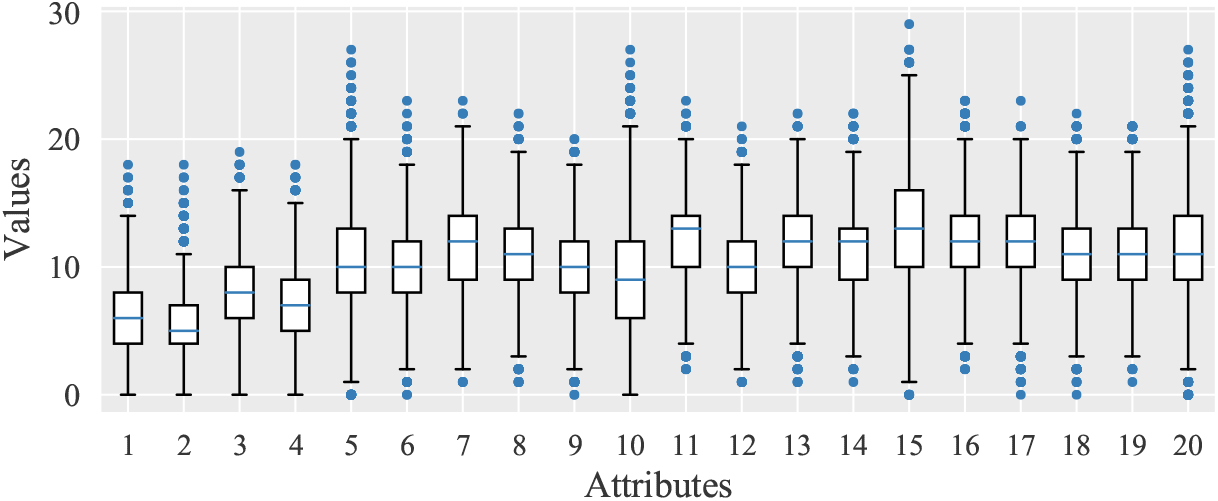
Sparsity of the first 20 attributes of original Mismatch profile. The x-axis represent the variation of data in each attribute produced by 4^1^ (1-4) mismatchs count concatenated with 4^2^ (5-20) and y-axis the variation interval of count.

**Figure 3:**
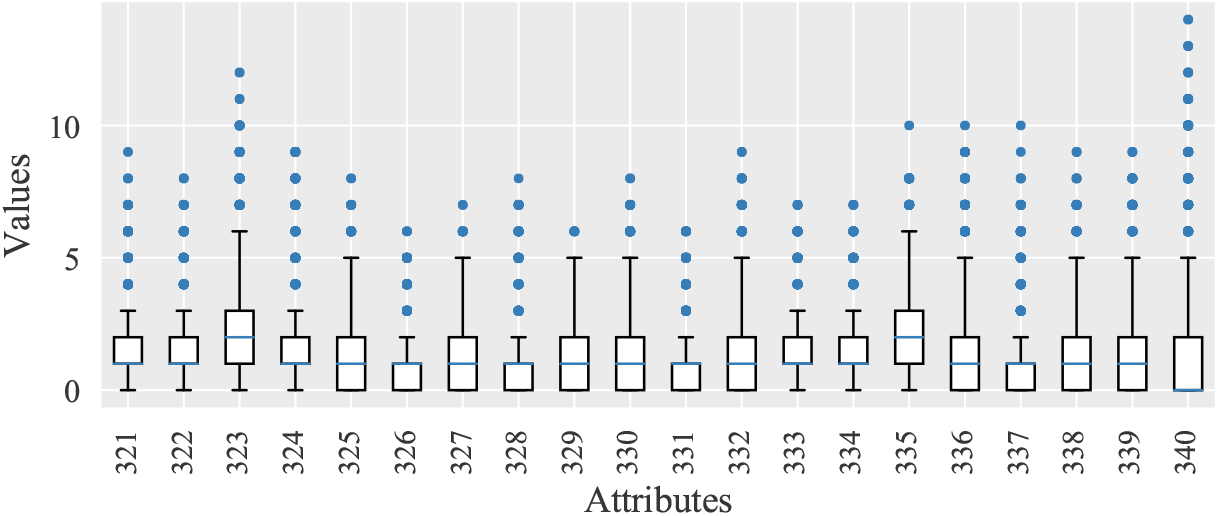
Sparsity of the last 20 attributes of original Mismatch profile, where x-axis represent the variation data in the last 20 attributes produced by 4^4^ (85-340) mismatchs count and y-axis the variation interval.

### 3.1. Analysis of Hyperparameter Optimization

Since each of the eight models were trained and evaluated in all nine datasets of group G1, a total of 72 results were obtained. In order to present the results in a simple and illustrative way, Figure 4 show the F-measures achieved by the models with 10-Fold Cross Validation in the hyperparameter optimization step. Note that the names of the models used for each configuration are defined in Table 1, where the X in DFNX stands for the number (identification) of the experiment performed (Deep Feedforward Network number X).

**Figure 4:**
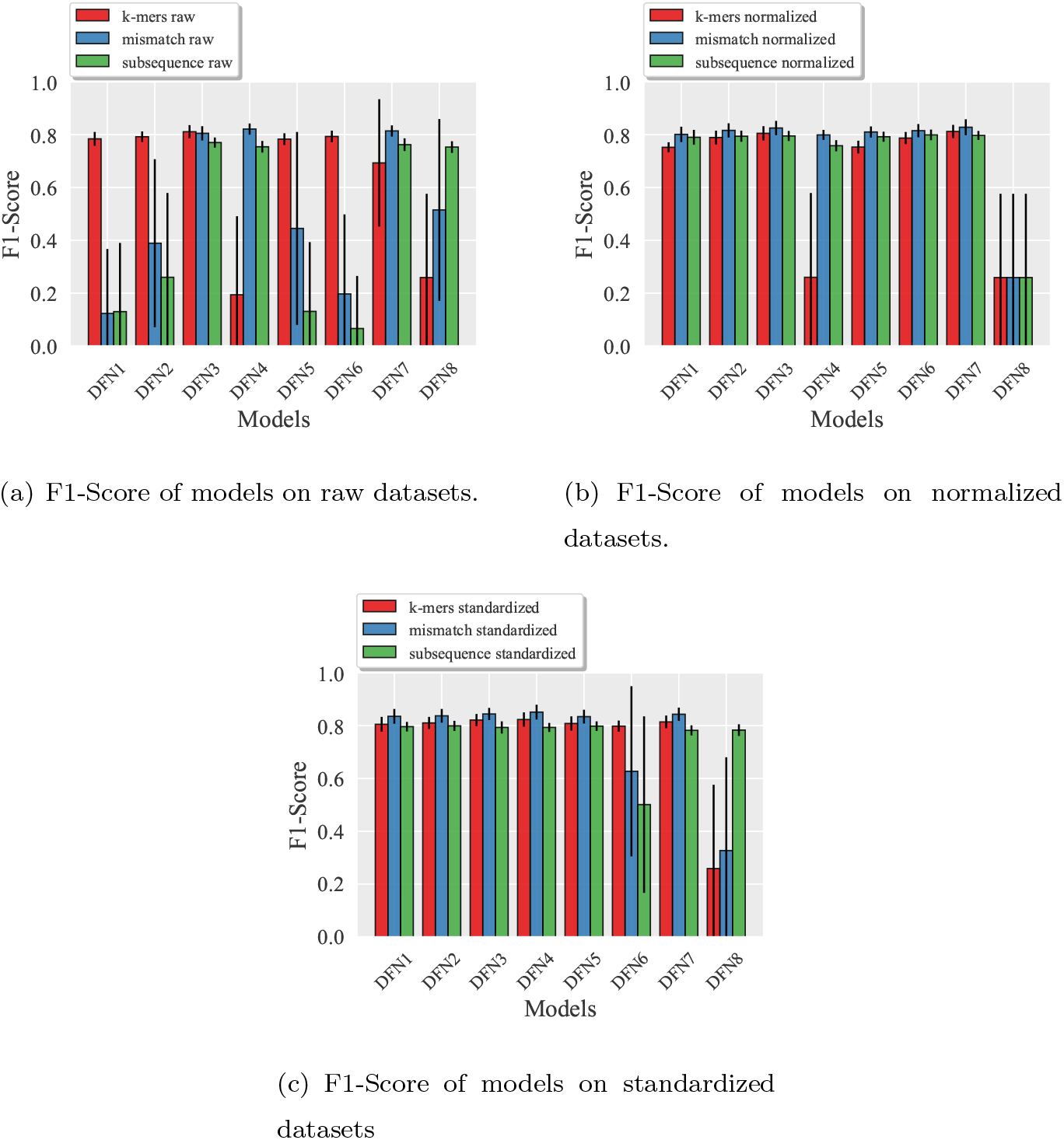
F1-Score of models on each dataset variant (i.e., raw, normalized and standardized). Each bar indicates the average achieved by the model in the K-Fold Cross Validation on that dataset (indicated by color in legend). The black lines on each bar indicates the standard deviation.

One important characteristic of deep learning models is the high computational power generated by the large number of hidden layers. However, from the results presented in Figure 4, we can see that increasing the depth of the neural networks did not improve their performance. Although DFN7 achieved the best result in *k*-mers and mismatch (both normalized) (with the same being observed for DFN6 in normalized subsequence), the differences between the results obtained compared with the three hidden layer neural networks are irrelevant. Even models with ReLU activation, which allows neural networks with a large number of hidden layers, did not have better results. Thus, it is observed that in the case of piRNAs prediction (and possibly other short ncRNAs), very deep models may not produce good results, even with an activation function adequate for deep models. It is also worth mentioning that the excess of complexity of a model can lead to the occurrence of overfitting [28], besides an unnecessary high computational cost.

Regarding the activation functions, ReLU has not been effective in producing better results in deeper neural networks, but obtained good results in the pre-processed datasets, even on standardized datasets (whose presence of negative values is significant) and without any vanishing gradient problem, considering that the increase of layers did not harmed the performance. However, it is performance on raw mismatch and subsequence were very poor, regardless of the number of layers or optimizer used, as can be seen in Figure 4(a). Note also from Figure 4(c) that DFN6 had a poor performance on standard mismatch and subsequence.

In contrast, some models with sigmoid had the performance impaired by increasing the number of hidden layers, mainly on raw and normalized *k*-mers. For example, DFN4 and DFN8 on raw *k*-mers (Figure 4(a)) performed very poorly and unstable, while DFN7 presented some instability. The only model with sigmoid and good results was DFN3. Moreover, DFN8 is the worst model, with a poor performance in seven of nine datasets, as shown in Figure 4. However, although some models with sigmoid have obtained poor results on some datasets, on raw datasets the performance was as good as the best results obtained on preprocessed data.

Thus, considering the results so far, the poor performances obtained by the neural nets with sigmoid must have occurred due to the vanishing gradient, since models with 3 layers (and sigmoid) had good results in general, but 5 layers model not. In addition, is known in the literature that neural networks with a large number of hidden layers together with sigmoid activation tend to have such a problem. At the same time, the properties that make sigmoid incapable of being used in many layers make it very powerful for dealing with the immense sparsity of the data, outliers and any other problem in the raw data, since models with sigmoid achieved good results on raw datasets and the DFN3 achieved good results in all datasets.

On other hand, ReLU had no vanishing gradient problems, but the sparse activation of ReLU may have caused the exploding gradient problem, since models with ReLU (mainly with SGD) were unable to learn from training data of raw mismatch and subsequence, whose spartisy is large, which can lead to very large gradient and, consequently, large updates to the network weights, producing an unstable network.

The optimization algorithms chosen also have significant impacts on the analyzed models. From Figure 4, we can see that several models with Stochastic Gradient Descent (SGD) failed to successfully execute the classification task due to the data or poor choice of hyperparameters. For example, DFN4 was unable to learn from both raw *k*-mers and normalized *k*-mers, and DFN8 was unable to learn from practically all datasets. On the other hand, DFN3 and DFN7, whose number of layers and activation function used correspond to same ones used in DFN4 and DFN8, reached a great performance on all datasets. Note that Adam was a better choice not only for models with sigmoid, but also models with ReLU. After all, since standardized datasets contain negative values, many neurons tend to be inactive (i.e., only 0 outputs), preventing learning from occurring. Thus, comparing the results obtained by DFN5 and DFN6 on standardized datasets, it is clear that the use of an optimization algorithm such as Adam is much more indicated in this case than the SGD. Adam is much more efficient to deal with noisy data or outliers, sparse gradients and bad hyperparameter choices, as can be seen in our experimental results.

### 3.2. Comparison with other methods

The best performing model in the hyperparameter optimization step was DFN3, which has three hidden layers, sigmoid activation and Adam optimizer. Considering that the best performance of DFN3 was on standardized mismatch (i.e., the mismatch profile rescaled by Z-Score Normalization), only these features were used in the comparison with other literature methods.

Our main rival state-of-the-art method is piRNN [14], since it is so far the only deep learning predictor for piRNAs. The piRNN method is a Convolutional Neural Network (CNN) where each sequence is a 4*×*341 matrix and each element represents *k*-mer motif frequencies. The first two layers are convolutional layers with a total of 32 2 *×* 2 filters, followed by a max pooling layer with 2 *×* 2 pooling size, and a dropout layer with 0.25 dropout ratio connected to the max pooling layer. The last part of the neural net is a dense layer with 512 nodes, a dropout layer with 0.5 dropout ratio, and the output layer with two nodes whose outputs correspond piRNAs (positive output) and non-piRNAs (negative output).

Besides piRNN, we also implement, train and evaluate two powerful traditional machine learning algorithms: Support Vector Machine (SVM) and Random Forest (RF). Both methods underwent a process of hyperparameter optimization (just like DFNs), on exactly the same datasets as the DFNs. The best configuration for SVM was *C* = 7.0, *γ* = 0.0005 and radial basis function kernel (where *C* is the penalty parameter and *γ* is the kernel coefficient gamma). For RF, we used 500 trees and entropy criterion.

The results obtained by all methods in 10-Fold Cross-Validation and G2 group samples are shown in Table 2. To compare the computational cost between DFN3 and piRNN, Table 2 also shows the total number of trainable parameters (Total params column) in both neural networks.

**Table 2:**
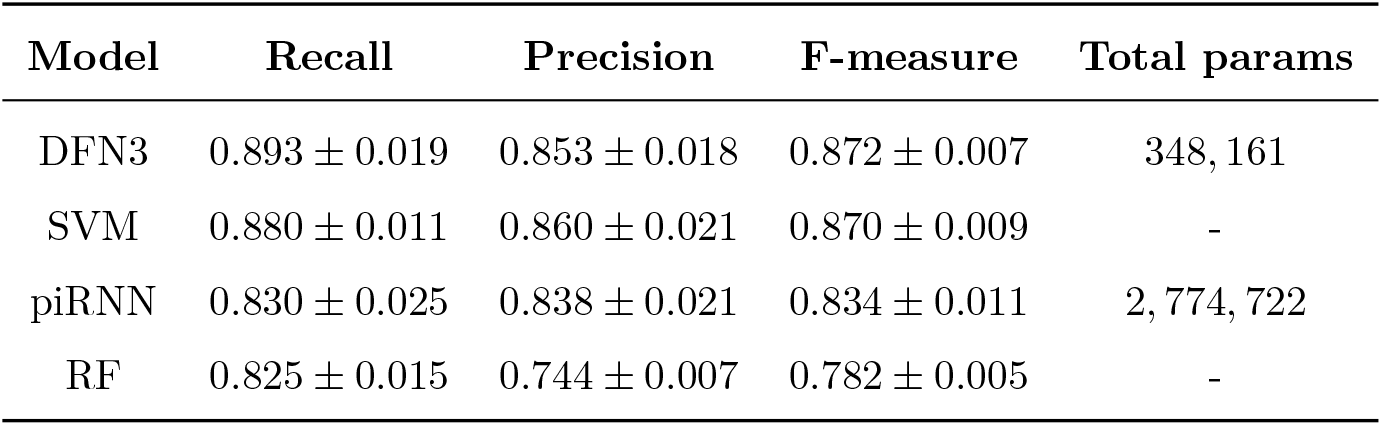
Comparison between DFN3, piRNN, SVM and RF

| Model | Recall | Precision | F-measure | Total params |
| --- | --- | --- | --- | --- |
| DFN3 | $0.893 \pm 0.019$ | $0.853 \pm 0.018$ | $0.872 \pm 0.007$ | 348,161 |
| SVM | $0.880 \pm 0.011$ | $0.860 \pm 0.021$ | $0.870 \pm 0.009$ | - |
| piRNN | $0.830 \pm 0.025$ | $0.838 \pm 0.021$ | $0.834 \pm 0.011$ | 2,774,722 |
| RF | $0.825 \pm 0.015$ | $0.744 \pm 0.007$ | $0.782 \pm 0.005$ | - |

From the results obtained by all predictors, we can see that our proposed model outperformed piRNN in all evaluation measures, specially Recall. The computational cost of our method is also much lower than the piRNN with a total number of parameters approximately 8 times smaller. Regarding the SVM and RF, SVM achieved excellent performance with better a Precision than our method, but Recall and F-measure were lower. RF although with a Recall close to the piRNN, it was the worst performing predictor. Thus, it is clear that despite the success and good performance of CNNs in classification tasks and their wide use in computational biology, DFNs can perform such prediction tasks as well as better than CNNs, achieving good results with less computational resources.

## 4. Conclusions

A Deep Feedforward Network is a basic architecture, but powerful and capable of successfully perform classification tasks in computational biology, including piRNAs prediction. Although very deep architectures have high computational power, they did not necessarily achieve excellent results. Thus, it is very important to correctly fit the number of hidden layers since a much more complex model than the problem can overfitting with an unnecessary high computational cost. ReLU activation functions, although being the state-of-the-art in avoiding the vanishing gradient problem, are not a good choice when the data have a large sparsity and many outliers, which is common for piRNA sequences and other ncRNAs (negative samples). Thus, the application of a feature scaling algorithm is essential for the use of ReLUs. On the other hand, sigmoid activation is very susceptible to the occurrence of vanishing gradient problems. However, it was very efficient to deal with the sparsity and outliers of our used data, reaching great results in the features before and after the feature scaling. Therefore, for both piRNAs and other ncRNAs, the use of sigmoid in DFNs may be a good solution. The correct choice of the optimization algorithm also has a significant impact on the neural network performance, with Adam being a better choice than the SGD for the data in question.

Finally, our model has achieved a great performance in piRNAs classification, outperforming the piRNN method, the state-of-the-art DNN for piRNAs classification. Thus, we showed that DFNs can be classifiers as good as better than CNNs. As future works, activation functions like LeakyReLU, ELU and Swish, and optimizers like Nadam and AMSGrad should be tested. Finally, we plan to extend our study and proposed model to other small ncRNAs, such as miRNAs, siRNAs and piRNAs (not only transposon-derived). This certainly can help computational biologists to build models with high classification performances.

